# A forward genetic screen identifies Dolk as a regulator of startle magnitude through the potassium channel subunit Kv1.1

**DOI:** 10.1101/2020.06.19.161240

**Authors:** Joy H. Meserve, Jessica C. Nelson, Kurt C. Marsden, Jerry Hsu, Fabio A. Echeverry, Roshan A. Jain, Marc A. Wolman, Alberto E. Pereda, Michael Granato

## Abstract

The acoustic startle response is an evolutionary conserved avoidance behavior. Disruptions in startle behavior, in particular startle magnitude, are a hallmark of several human neurological disorders. While the neural circuitry underlying startle behavior has been studied extensively, the repertoire of genes and genetic pathways that regulate this locomotor behavior has not been explored using an unbiased genetic approach. To identify such genes, we took advantage of the stereotypic startle behavior in zebrafish larvae and performed a forward genetic screen coupled with whole genome analysis. This identified mutants in eight genes critical for startle behavior, including two genes encoding proteins associated with human neurological disorders, Dolichol kinase (Dolk), a broadly expressed regulator of the glycoprotein biosynthesis pathway, and the potassium Shaker-like channel subunit Kv1.1. We demonstrate that Kv1.1 acts independently of supraspinal inputs to regulate locomotion, suggesting its site of action is within spinal circuitry. Moreover, we show that Kv1.1 protein is mis-localized in *dolk* mutants, suggesting they act in a common genetic pathway to regulate movement magnitude. Combined, our results identify a diverse set of eight genes all associated with human disorders that regulate zebrafish startle behavior and reveal a previously unappreciated role for Dolk and Kv1.1 in regulating movement magnitude via a common genetic pathway.

**Author summary:** Underlying all animal behaviors are neural circuits, which are controlled by numerous molecular pathways that direct neuron development and activity. To identify and study these molecular pathways that control behavior, we use a simple vertebrate behavior, the acoustic startle response, in the larval zebrafish. In response to an intense noise, larval zebrafish will quickly turn and swim away to escape. From a genetic screen, we have identified a number of mutants that behave in abnormal ways in response to an acoustic stimulus. We cloned these mutants and identified eight genes that regulate startle behavior. All eight genes are associated with human disorders, and here we focus on two genes, *dolk* and *kcna1a*, encoding Dolk, a key regulator of protein glycosylation, and the potassium channel Kv1.1, respectively. We demonstrate that loss of *dolk* or *kcna1a* causes larval zebrafish to perform exaggerated swim movements and that Dolk is required for Kv1.1 protein localization to axons of neurons throughout the nervous system, providing strong evidence that *dolk* and *kcna1a* act in a common molecular pathway. Combined, our studies provide new insights into the genetic regulation of startle behavior.

## Introduction

Defects with initiating or executing movements are associated with a range of disorders. While some neurological disorders are primarily defined by motor impairments, several disorders defined primarily by cognitive deficits include motor features. For example, the eyeblink response, in which a patient reacts to a startling stimulus, is disrupted in a variety of neurodevelopmental and psychiatric disorders, including obsessive compulsive disorder, schizophrenia, posttraumatic stress disorder, and autism spectrum disorder [1-4]. The eyeblink response is one component of the startle response, which in humans is a whole-body defensive maneuver to shield the upper body from impact and in aquatic vertebrates, including zebrafish, is critical to evade avoid predators [5,6]. A combination of electrophysiological, lesion and imaging studies have uncovered the core neural circuity underlying startle behavior in human and various vertebrate animal models [7-10]. Yet despite its critical role in animal survival and its link to several human neurological disorders, the repertoire of genes and genetic pathways that regulate startle behavior has not been explored using an unbiased genetic approach.

Over the past several decades, the larval zebrafish has emerged as a powerful vertebrate model organism for unbiased genetic screens to identify genes critical for basic locomotion [11-13] and more recently for more complex behaviors, including visual behaviors and sleep [14,15]. However, an unbiased genetic screen to identify the genes critical for the execution of the startle response has been absent. By five days post fertilization (dpf), in response to an acoustic stimulus, zebrafish larvae undergo a characteristic short latency C-start (SLC), consisting of a sharp C-shaped turn and swimming away from the stimulus [16] (Fig 1B-F, Video S1). The behavioral circuit for the acoustic startle response is well characterized (reviewed in [9,17], Fig 1A) and is functionally similar to the human startle circuit [5]. Central to the zebrafish acoustic startle circuit are the Mauthner cells, a bilateral pair of reticulospinal neurons in the hindbrain. The Mauthner cells are necessary and sufficient for this short latency escape behavior [16,18,19]. Hair cell activation following an acoustic stimulus leads to activity of the eighth cranial nerve, which, along with the spiral fiber neurons, provides excitatory input to the Mauthner cell. The Mauthner cell directly activates contralateral primary motor neurons and excitatory interneurons to drive unilateral body contraction and turning away from the acoustic stimulus [20,21]. Inhibitory input prevents Mauthner cell firing at subthreshold stimulus or when the other Mauthner cell has already fired [22]. Because the circuit is well defined and the behavior is robust, this system is ideal for investigating how genes regulate behavior.

**Fig 1.**
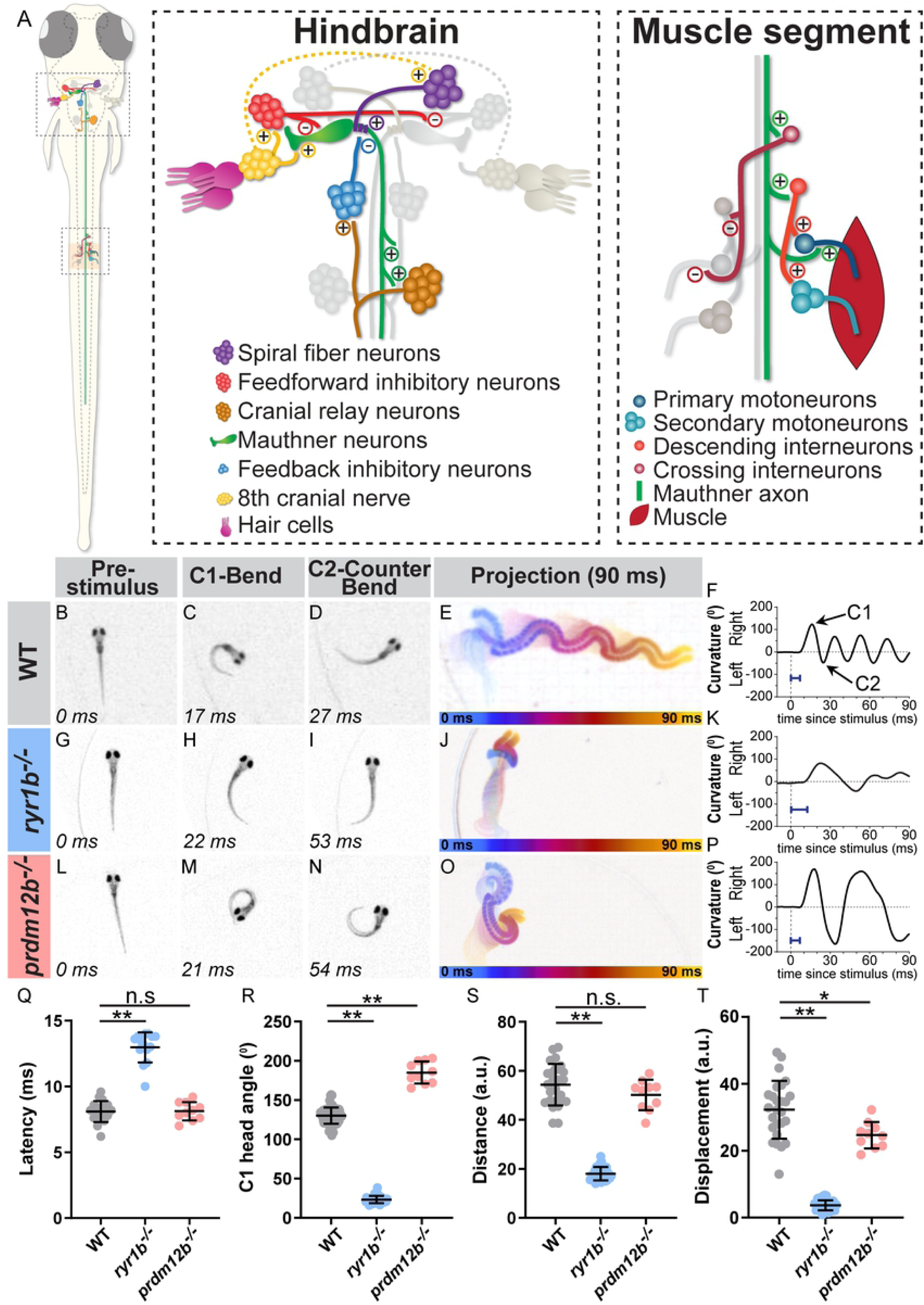
The zebrafish larval startle response is amenable to circuit and genetic analysis. (A) The acoustic startle response is driven by an action potential from the Mauthner neuron (green), which synapses on motor neurons in the spinal cord to drive a contralateral body bend. Excitatory and inhibitory neurons in the hindbrain and in the spinal cord impinge upon the Mauthner cells to ensure motor neurons on only one side fire. (B-F) A representative acoustic startle response in a 5 dpf wild type larva. An acoustic stimulus is delivered at 0 ms (B), which elicits a rapid turn (C) followed by a counter bend (D) and swimming away (projection of 90 ms response in E, color coded by time). Automated tracking of the curvature of the larva throughout the behavior (F) reveals the response latency (blue bar=latency in F,K,P). (G-K) *ryr1b* mutants display a weak startle response with reduced bend and counter bend angles (H,I) and reduced displacement (J), largely due to minimal swimming after the counter bend (K). (L-P) *prdm12b* mutants display an exaggerated startle response with increased bend and counter bend angles (M,N). The duration of each bend is longer than in wild type as well (P). (Q-T) Quantification of response latency (Q; manual measurement); max C1 head angle (R; automated measurement); distance traveled along escape trajectory (S; automated measurement); and displacement from initial head position to final head position (T; automated measurement). Each point is the average response over ten trials for an individual larva. n≥10 larvae, *p=0.001, **p<0.0001

We previously performed a forward genetic approach to identify functional regulators of the acoustic startle response. This approach identified a distinct sets of genes with previously unrecognized roles critical for startle habituation [23,24], startle sensitivity [25], and sensorimotor decision making [26]. Here, we describe kinematic mutants identified from this screen that display defects in executing swim movements of the startle response. These mutants fall into two general categories. One group of mutants display a “weak” acoustic startle response characterized by shallow bends and minimal displacement. Through genome wide sequencing, we have identified causative mutations in five weak startle mutants. In all five lines, the affected genes control muscle or neuromuscular junction (NMJ) function and are associated with locomotor disorders in humans.

The second group of mutants perform high amplitude bends, resulting in an “exaggerated” response. For one of the exaggerated startle mutant lines, we identified a causative mutation in *PR domain containing 12b* (*prdm12b*), a transcription factor that controls development of inhibitory neurons in the spinal cord and has been previously shown to be important for regulation of movement in fish [27]. The remaining two exaggerated mutant lines harbor mutations in *dolichol kinase* (*dolk*), which encodes a glycosylation pathway enzyme, and in *potassium voltage-gated channel, shaker-related subfamily, member 1a* (*kcna1a*), which encodes the potassium channel subunit Kv1.1. We demonstrate that *dolk* and *kcna1a* likely act in a common pathway to control startle movement magnitude as Dolk is required for Kv1.1 protein localization. Additionally, we demonstrate that Kv1.1 acts in the spinal cord to control the magnitude of body bends. Thus, our forward genetic screen identified a number of genes that are essential to regulate body movement. Furthermore, we demonstrate how a broadly expressed protein, Dolk, specifically affects behavior through regulation of a downstream neuronal protein, Kv1.1.

## Results

### A forward genetic screen for regulators of the larval startle response

We previously performed a forward genetic screen for regulators of the acoustic startle response [23-26]. In brief, using a high-speed camera and automatic tracking, individual F3 larvae were exposed to a series of startling stimuli (for details on mutagenesis and the breeding scheme to obtain F3 larvae see [23]). This portion of the screen focused on kinematic mutants, characterized by significant changes in any of the stereotypic parameters characteristic for the startle response (Table 1). Putative mutant lines were retested in the subsequent generation to confirm genetic inheritance. This screen identified a group of eight mutants defective in kinematic parameters (including response latency, turn angle, distance, and displacement). These mutants fell into two categories: five mutant lines display shallow bends of decreased turning angle compared to their siblings (one representative mutant line shown in in Fig 1G-K,Q-T, Video S2); we refer to these as “weak” startle mutants. Three mutant lines perform numerous high amplitude bends in response to an acoustic stimulus. These turns often result in larvae swimming in a figure eight pattern (one representative mutant line shown in Fig 1L-T, Video S3); we refer to these as “exaggerated” startle mutants. Combined, these eight mutant lines offer an opportunity to reveal genetic regulators of movement kinematics.

**Table 1.**
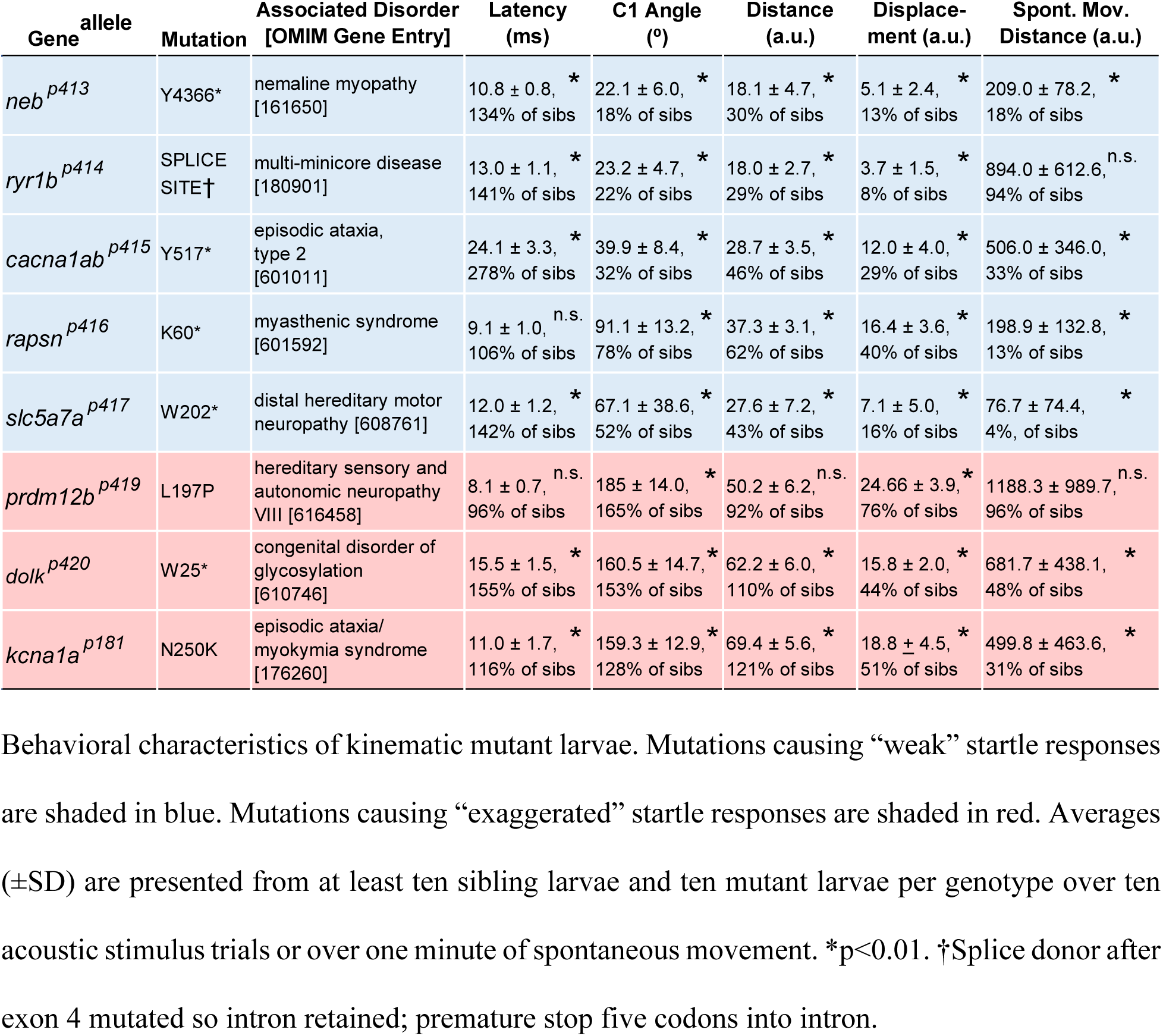
Genes identified from a forward genetic screen that regulate locomotor behaviors in larval zebrafish.

To identify the molecular mechanisms underlying these behavioral phenotypes, we first set out to identify the causative mutations. In all kinematic mutant lines, approximately 25% of progeny from carrier incrosses display the mutant behavioral phenotype, consistent with the causative mutations being recessive, monoallelic, and causing highly penetrant phenotypes. For each mutant line, after behavioral testing, pools of behaviorally mutant larvae and behaviorally wild type siblings were collected for high-throughput DNA sequencing. We performed whole genome sequencing (WGS) and homozygosity analysis on mutant and sibling pools for four of the lines, as described in more detail in [23]. Using known single nucleotide polymorphisms (SNPs) present in our wild type background, we identified regions of homozygosity in the mutant pools that were heterozygous in the sibling pool. Within that region, potentially detrimental exonic SNPs not observed in wild type fish were identified. Nonsense mutations, particularly in genes known to function in neurons or muscle, were prioritized. Individual larvae displaying mutant or wild type behavior were then sequenced for potentially causative SNPs. If a potential SNP was observed as homozygous in 100% of mutant larvae (>20 individuals) and heterozygous or homozygous wild type in all siblings (>20 individuals), we considered the SNP to likely be causative. For the remaining four mutant lines, we performed whole exome sequencing (WES) [28]. To identify linkage (SNPs with allelic frequencies ∼100% in mutants and ∼33% in siblings, based on ratio of heterozygous to homozygous wild type larvae) and potentially causative mutations, we used the online tool SNPTrack [29]. Confirmation of potentially causative mutations was performed as described above for candidates from WGS. Five lines (*neb, cacna1ab, ryr1b, rapsyn*, and *slc5a7a*) contain nonsense mutations (or a splice mutation resulting in nonsense mutation) in genes known to regulate neuron or muscle function (see Table 1), strongly indicating we have identified the correct mutation (see also discussion of identified genes below). Two of the mutant lines (*prdm12b* and *kcna1a*) have missense mutations, and we or others have made nonsense alleles that display the same phenotype. For the remaining mutant line (*dolk*) containing a nonsense mutation, we have generated an independent second mutant allele to confirm that mutations in *dolk* are causative for the exaggerated locomotor phenotype (see below). Based on these data, we are confident we have identified mutant alleles of eight genes critical for regulating proper kinematic behavior during the acoustic startle response. We note that all genes have a human disease associated counterpart and that six of the eight genes are associated with human movement disorders, further underscoring conservation of disease associated genes in zebrafish [30] (Table 1). Based on the molecular identity, the affected genes can be subdivided by their likely site of action. Below, we report on several genes that act in skeletal muscle, at the neuromuscular junction, or in inhibitory spinal neurons, and we report in detail on two genes that appear to act in the same pathway to regulate movement magnitude.

### Genes controlling muscle function modulate acoustic startle movement kinematics

Of the five weak startle mutant lines, two have mutations in genes that are required for muscle function. *p413* mutants contain a nonsense mutation in the *nebulin* (*neb*) gene (Table 1), which encodes a protein necessary for sarcomere assembly and subsequent function [31]. Previous work in zebrafish demonstrated that a loss-of-function *neb* allele displays defects in sarcomere assembly, leading to reduced swim movement [32]. The second mutant line we identified, *p414*, contains a splice mutation in the *ryanodine receptor 1b* (*ryr1b*) gene (Table 1; Fig 1G-K,Q-T). RyR1 is required for calcium release at the sarcoplasmic reticulum, which drives muscle contraction. A previously characterized zebrafish allele of *ryr1b*, called *relatively relaxed*, displays a decreased touch response and has reduced Ca^2+^ transients in fast muscle [33]. Interestingly, RyR1b is expressed primarily in fast muscle while RyR1a is expressed primarily in slow muscle [33]. Consistent with this finding, *ryr1b*^*p414*^ mutants display a drastic startle response defect, which is dependent on fast muscle. However, spontaneous movement, which is dependent on slow muscle, was not significantly different from siblings (Table 1). This result emphasizes the importance of examining different behaviors, especially ones that utilize different circuitry and muscle groups [20,34,35], when characterizing locomotion in mutant animals.

### Genes acting at the neuromuscular junction regulate acoustic startle movement kinematics

Three genes identified in our screen are required at the neuromuscular junction (NMJ). This includes a new mutant allele, *p416*, of the gene *receptor-associated protein of the synapse* (*rapsyn*), originally identified as *twitch once* mutants [11]. *rapsyn* mutants display reduced spontaneous swimming and touch response [11] caused by reduced acetylcholine receptor clustering at NMJs [36]. In addition, we identified a nonsense mutation (*p415*) in the *calcium channel, voltage-dependent, P/Q type, alpha 1A subunit, b* (*cacna1ab*) gene, which encodes Ca_v_2.1. A previously identified mutant allele of *cacna1ab, fakir*, displays a weak touch response [11,37]. This weak touch response in *fakir* larvae can be attributed to defects in transmission at the NMJ [38]. This can account for the reduction in startle response and spontaneous movement we observe in *p415* (Table 1). Since *rapsyn* and *cacna1ab* were both previously identified in a screen for escape response following a tail touch [11], which is driven by the Mauthner cells [39], we predicted them to be identified in the Mauthner-dependent acoustic startle screen described here.

The fifth weak startle mutant, *p417*, demonstrates a unique behavior (Fig S1). Unlike *p417* sibling larvae that initiate the startle response at the head (Fig S1C-G), in *p417* mutants, the startle response often initiates with a turn in the tail instead (Fig S1H-L, Video S4). Subsequent body bends are uncoordinated, rather than a smooth progression from head to tail. The C1 angle, distance, and displacement are also drastically reduced in these mutants, and mutants undergo very little spontaneous movement (Table 1). We identified the causative *p417* mutation as a nonsense mutation in *slc5a7a*, which encodes the high-affinity choline transporter (CHT) (Fig S1A-B). Uptake of choline by the high-affinity choline transporter is the rate limiting step for acetylcholine synthesis [40], so Slc5a7 is essential for cholinergic transmission.

There are a number of cholinergic neuronal populations in the acoustic startle circuit. Most notably, spinal motor neurons release acetylcholine at the NMJ to activate skeletal muscle in the trunk and tail [41]. We hypothesized the *slc5a7a* mutant phenotype of reduced turn angle and minimal displacement is caused by reduced muscle contractions due to reduced acetylcholine release at the NMJ. To test this, we generated a transgenic line in which *slc5a7a* is expressed under the control of a motor neuron specific promoter, *mnx1/hb9* [42] (Fig S1M). Compared to *slc5a7a* mutants lacking the transgene (Fig S1N-P), in mutant larvae expressing the transgene (*slc5a7a*^*-/-*^; *Tg(hb9:slc5a7a)*, the turn angle and distance traveled following an acoustic stimulus is significantly increased. Thus, motor neuron specific expression of *slc5a7a* partially rescues the mutant phenotype, providing strong evidence that Slc5a7a plays a critical role in regulating coordinated swim movements, presumably by selectively increasing acetylcholine level release at NMJs.

### *prdm12b* regulates development of V1 interneurons and regulates acoustic startle movement kinematics

In addition to the five weak startle mutants described above, we identified three exaggerated startle mutants. In the *p419* mutant line in which the C1 turning angle of the startle response is dramatically increased (165% of wild type siblings; Table 1; Fig 1L-P, Video S3), we identified a missense mutation in the coding sequence of *PR domain containing 12b* (*prdm12b*). Prdm proteins are transcription factors, and several family members have been shown to play roles in nervous system development [43]. Zebrafish *prdm12b* was previously identified in a screen for genes that express in the nervous system [44]. Morpholinos targeting *prdm12b* [27] and a *prdm12b* CRISPR mutation [45] result in larvae with aberrant, exaggerated touch behaviors, similar to the exaggerated acoustic startle response we observe in the *p419* mutants. *prdm12b* is required for development of *engrailed 1b* positive, V1 interneurons in the spinal cord [27]. V1 interneurons are glycinergic and have been shown to regulate the speed of locomotor movements in mammals [46]. Thus, Prdm12b is required during development for the specification of inhibitory spinal neurons utilized for the acoustic startle response to achieve coordinated and controlled movement.

### The glycosylation pathway protein Dolk regulates swim magnitude

In a second exaggerated movement mutant, *p420*, we identified a nonsense mutation in the *dolichol kinase* (*dolk*) gene (Fig 2A,B). Similar to *prdm12b* mutants, the *dolk* mutant C1 turning angle is dramatically increased (153% of wild type siblings; Table 1; Fig 2C-L,M). In addition to exaggerated turn angles in response to an acoustic stimuli, *dolk* mutants swim in “figure eights,” resulting in reduced displacement (Fig 2K,O). Dolk catalyzes the CTP-dependent phosphorylation of dolichol to dolichol monophosphate. Dolichol monophosphate is an essential glycosyl carrier for C- and O-mannosylation and N-glycosylation of proteins [47]. To confirm that the *dolk* nonsense mutation identified in our screen (*p420*) is causative, we generated a second mutant *dolk* allele using CRISPR/Cas9. In this *dolk* ^*p421*^ CRISPR allele, a 1 bp deletion results in a frameshift and premature stop codon at position 50. Transheterozygous (*p420/p421*) *dolk* mutants display the same exaggerated phenotype as *p420* homozygotes (Fig 2M-O), confirming that mutations in *dolk* cause the behavioral deficits observed in the original *p420* mutants.

**Fig 2.**
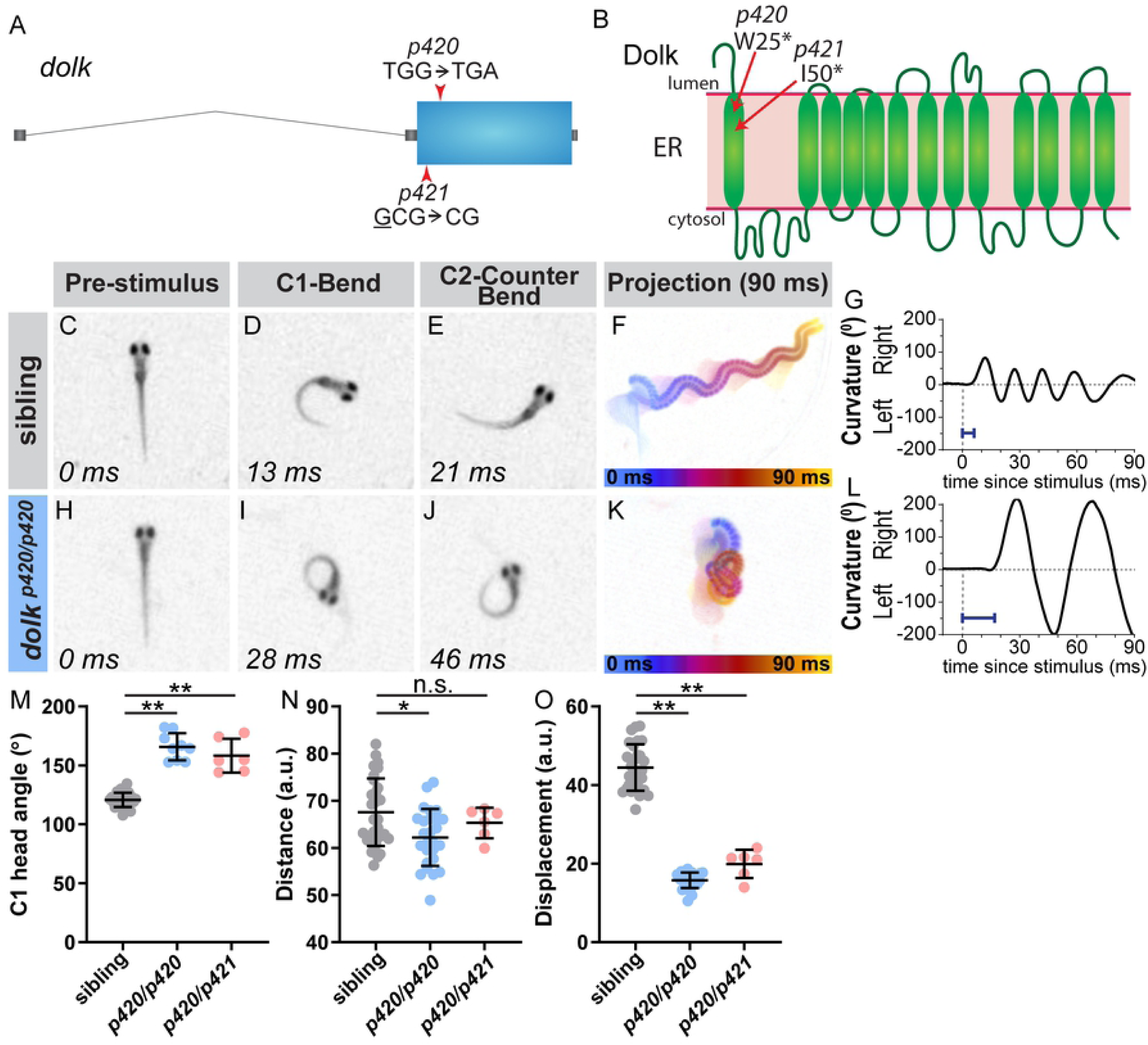
The glycosylation pathway enzyme Dolk regulates the magnitude of the startle response. (A) Gene structure for *dolk* with the nonsense mutation from the screen noted (*p420*). The CRISPR 1bp deletion allele is also noted (*p421*). (B) Protein structure for Dolk on the endoplasmic reticulum, with the predicted amino acid change from the screen mutation indicated. The deletion in the CRISPR allele causes a frame shift in the sequence that results in a premature stop at amino acid 50. In contrast to siblings (C-G), *dolk* mutants (H-L) display an exaggerated startle response, with an increased bend and counter bend angle (I,J), resulting in larvae swimming in a “figure eight” (K). Blue bar=latency in G,L. (M-O) Kinematic parameters of the acoustic startle response in *dolk* siblings and mutants (homozygotes of the screen identified mutation, *p420/p420*, and transheterozygotes from the screen mutation and CRISPR, *p420/p421*). Each point represents average of ten trials for an individual fish. n≥6 larvae, **p<0.001, *p<0.01

Given Dolk’s critical role in protein glycosylation, we predict that in *dolk* mutants, many proteins exhibit reduced glycosylation. In 2 dpf zebrafish, over 160 unique glycosylated proteins have previously been cataloged via mass spectrometry [48], so predicting the number and identities of Dolk’s behavior-relevant glycosylation targets is challenging. To identify the relevant glycosylated downstream protein(s), we instead turned to our remaining exaggerated mutant line. In this line, we identified a missense mutation in *kcna1a*, which encodes the voltage potassium Shaker-like channel subunit Kv1.1 (Fig 3A,B) [24]. Cell culture studies have shown that Kv1.1 is highly glycosylated, and glycosylation is required for Kv1.1 channel function [49-51]. We confirmed the kinematic defect in the screen *p181* allele (Fig 3C-O) is due to the missense mutation in *kcna1a* using a previously generated *kcna1a* CRISPR allele (*p410*) [24]. *p181/p410* transheterozygotes display the same exaggerated movements as mutants homozygous for *p181* (Fig 3M-O). Thus, *kcna1a* and *dolk* are both required for executing controlled movements, possibly via Dolk-dependent glycosylation of Kv1.1.

**Fig 3.**
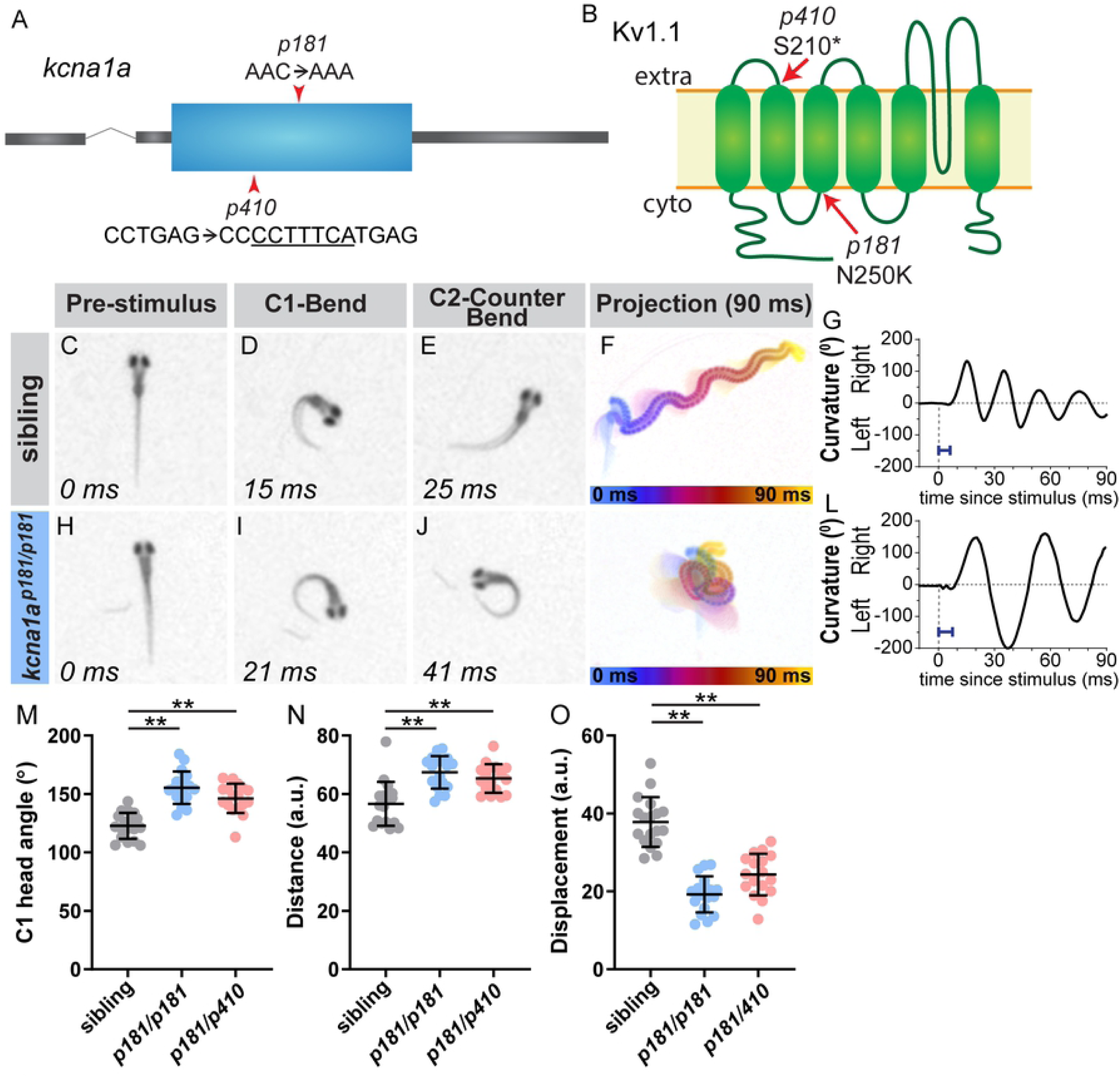
The potassium channel subunit Kv1.1 regulates the magnitude of the startle response. (A) Gene structure for *kcna1a* with the missense mutation from the screen noted (*p181*). The CRISPR 7 bp insertion allele is also noted (*p410*). (B) Protein structure for Kv1.1, which is encoded by *kcna1a*, on the plasma membrane, with the predicted amino acid change from the screen allele (*p181*) indicated. The insertion in the CRISPR allele (*p410*) causes a frame shift in the sequence that results in a premature stop at amino acid 210. In contrast to siblings (C-G), *kcna1a* mutants (H-L) display an exaggerated startle response, with an increased bend and counter bend angle (I,J), resulting in fish swimming in a “figure eight” (K). Blue bar=latency in G,L. (M-O) Kinematic parameters of the acoustic startle response in *kcna1a* sibling and mutants (homozygotes of the screen mutation, *p181/p181*, and transheterozygotes from the screen mutation and CRISPR, *p181/p410*). Each point represents average of ten trials for an individual fish. n≥17 larvae, **p<0.001

### Dolk is required for axonal localization of Kv1.1

Glycosylation is well known to regulate protein activity, localization, and/or stability [52]. To address whether *dolk* is critical for Kv1.1 localization or stability, we examined Kv1.1 protein localization in *dolk* mutants using a previously validated antibody [24]. In mammals, Kv1.1 is broadly expressed throughout the brain and primarily localizes to axons and axon terminals [53]. In 5 dpf wild type larvae, we observe Kv1.1 protein localization to axon tracts throughout the brain (Fig 4A-B) and the spinal cord (Fig 4C). Kv1.1 strongly accumulates at the Mauthner axon cap (Fig 4B), where spiral fiber and inhibitory neurons synapse onto the Mauthner axon initial segment (AIS). In contrast, in *dolk* mutants, Kv1.1 protein localization is strongly reduced or absent in axon tracts and instead is prominent in somata throughout the brain (Fig 4D-F). This is particularly apparent in the large Mauthner soma (Fig 4E-F). Little or no Kv1.1 is detectable at the axon cap (Fig 4E-F) in *dolk* mutants. Since we do still observe Kv1.1 expression in *dolk* mutants, Dolk may be largely dispensable for Kv1.1 protein stability yet indispensable for correct Kv1.1 cell surface localization. As previous studies have shown that Kv1.1 membrane localization is required for its channel function, the most parsimonious explanation for the behavioral phenotype in *dolk* mutants is the lack of proper Kv1.1 localization.

**Fig 4.**
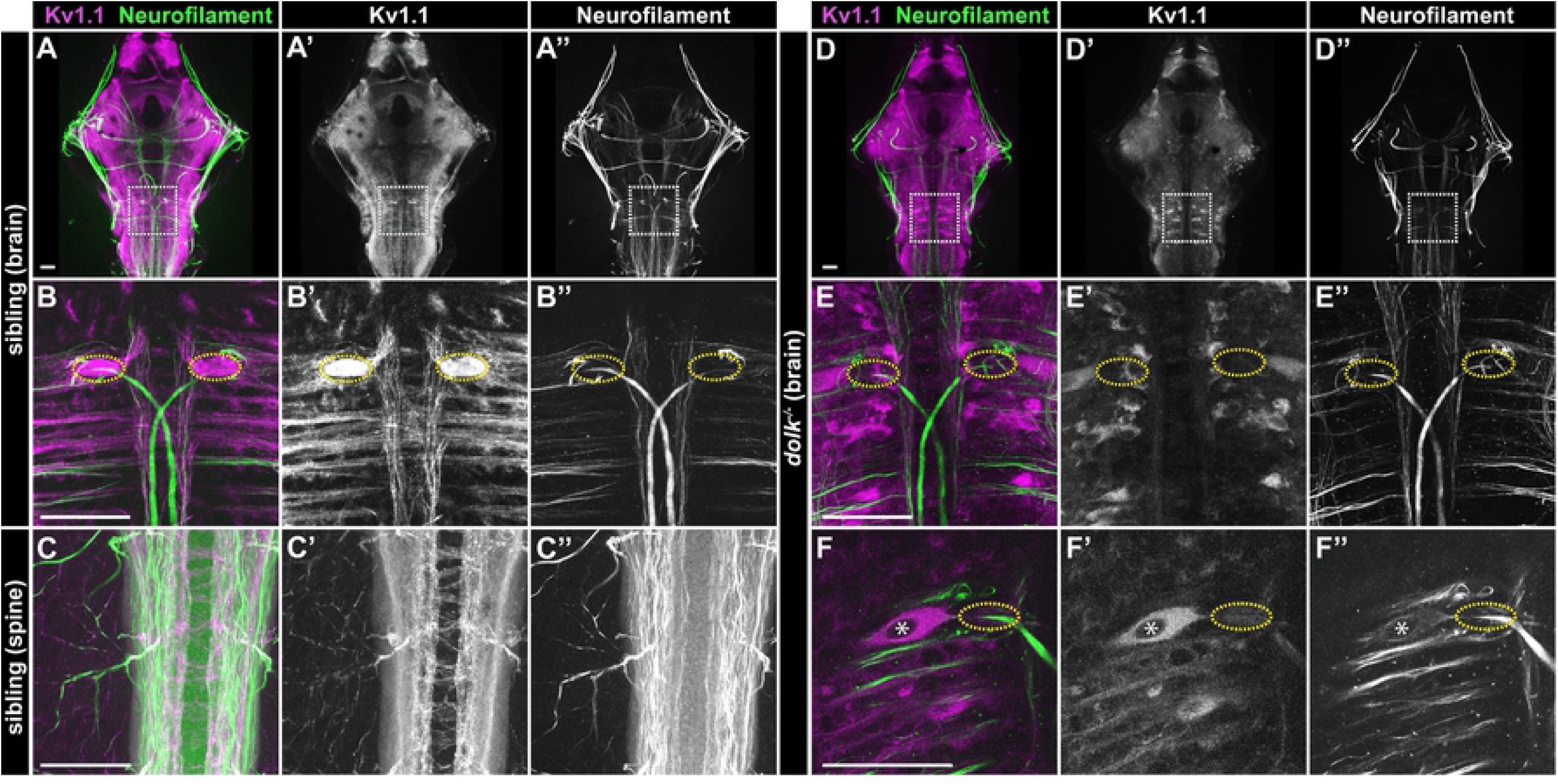
Dolk is required for proper localization of Kv1.1. (A-C) In 5 dpf *dolk* sibling larvae, Kv1.1 (magenta:A,B,C; grey:A’,B’,C’) localizes to fiber tracts (α-3A10 antibody stains neurofilament; green:A,B,C; grey:A”,B”,C”) throughout the brain (A; box indicates hindbrain zoom in B) and spinal cord (C). Particularly high accumulation of Kv1.1 is observed at the axon cap (B, yellow dotted circle), where spiral fiber neurons form axo-axonic synapses with the Mauthner cell. (D-F) In *dolk* mutants, Kv1.1 is localized within somata of neurons throughout the brain and is strongly reduced along axons throughout the high brain (D,D’,E,E’). Soma localization is particularly apparent in the large Mauthner neuron and strikingly absent from its axon cap (F, nucleus marked by asterisk, E,F axon cap marked by dotted ovals). Scale bars=50 µM.

### Kv1.1 acts in the spinal cord to regulate swim movement magnitude

We next asked where in the startle circuit Kv1.1 acts to regulate the magnitude of the startle response. Expression of Kv1.1 throughout the hindbrain and spinal cord is consistent with the idea that Kv1.1 may act at various sites in the startle circuit. *kcna1a* mRNA is observed in Mauthner neurons as early as 2 dpf, and developmental expression of Kv1.1 correlates with the Mauthner cell acquiring single spike properties [54,55]. Furthermore, inhibition of Kv1 channels by dendrotoxin-I injection into the Mauthner cell lowers the threshold required to induce multiple spikes in the Mauthner cell [54,55]. Based on these previous findings, it is possible that *kcna1a* functions in Mauthner neurons to regulate the magnitude of the startle response. We explored this possibility using whole-cell recordings to determine the electrophysiological properties of Mauthner cells in *kcna1a* mutant and sibling larvae. Averaged measurements of membrane resting potential (V_resting_), input resistance (R_in_), rheobase (amount of current necessary to trigger an action potential), and action potential threshold (V_threshold_) are not significantly different between *kcna1a* siblings and *kcna1a* mutant larvae (Fig S2A). Next, we asked whether loss of *kcna1a* function changes the weight of key synaptic inputs to the Mauthner cells. Auditory afferents known as “large myelinated club endings” (club endings) that terminate on the lateral dendrite of the Mauthner cell are critical inputs for the acoustic startle response [56,57]. To test whether this synaptic input was altered, we examined mixed (electrical and chemical) synaptic potentials evoked by electrical stimulation of club endings in sibling and *kcna1a* mutant larvae during whole-cell recordings on the Mauthner cell (Fig S2B). We observed no difference in the average amplitudes of either the electrical (Fig S2C) or chemical (Fig S2D) components of the mixed synaptic response, nor in the paired-pulse ratio of the chemical component (Fig S2E) indicating no changes in presynaptic neurotransmitter release. Combined, these results reveal that neither changes of the excitability of Mauthner cells nor changes in the weight of critical auditory synaptic inputs can explain the exaggerated startle behavior observed in *kcna1a* mutants.

While Mauthner cell function and club endings appear unaffected in *kcna1a* mutants, it is possible that other inputs onto the Mauthner cells or synapses of the Mauthner cells are disrupted in *kcna1a* mutants. To determine whether *kcna1a* acts in a Mauthner-dependent pathway, we ablated Mauthner cells in *kcna1a* mutant larvae. Following Mauthner cell ablation, wild type larvae presented with an acoustic stimulus perform a longer latency, Mauthner-independent turning maneuver [16]. Thus, analyzing behavior in *kcna1a* mutants in which Mauthner neurons are ablated can clarify whether the exaggerated phenotype is dependent on Mauthner cell function and/or the function of neurons ‘upstream’ of Mauthner cells. We chemogenetically ablated Mauthners in *kcna1a* siblings and mutants using the nitroreductase (NTR)/metronidazole (Mtz) system [58]. Prior to Mtz treatment, larvae were scored for presence or absence of Mauthner cell NTR-RFP expression (Fig 5A,B). Following Mtz treatment, in larvae initially lacking Mauthner NTR-RFP expression, Mauthner neurons were intact (Fig 5C). Conversely, following Mtz treatment in larvae with initial Mauthner NTR-RFP expression, Mauthner neurons were undetectable (Fig 5D). We then presented fish with intact (controls) or ablated Mauthner cells with an acoustic stimulus. As expected, compared to controls, sibling and *kcna1a* mutant larvae in which Mauthner cells were ablated display a longer response latency characteristic of Mauthner-independent turning maneuvers (Fig 5E) [16]. Importantly, *kcna1a* mutant larvae in which Mauthner cells were ablated still performed exaggerated responses and figure eight turns (Fig 5F-H). These responses were, aside from latency, indistinguishable from responses in mutants with intact Mauthner cells (Video S5). Thus, the exaggerated behavioral response in *kcna1a* mutants is independent of activity in neurons ‘upstream’ of the Mauthner cell or in the Mauthner cell itself, further supporting the notion that Dolk and Kv1.1 primarily act outside the Mauthner-dependent hindbrain startle circuit to regulate startle magnitude.

**Fig 5.**
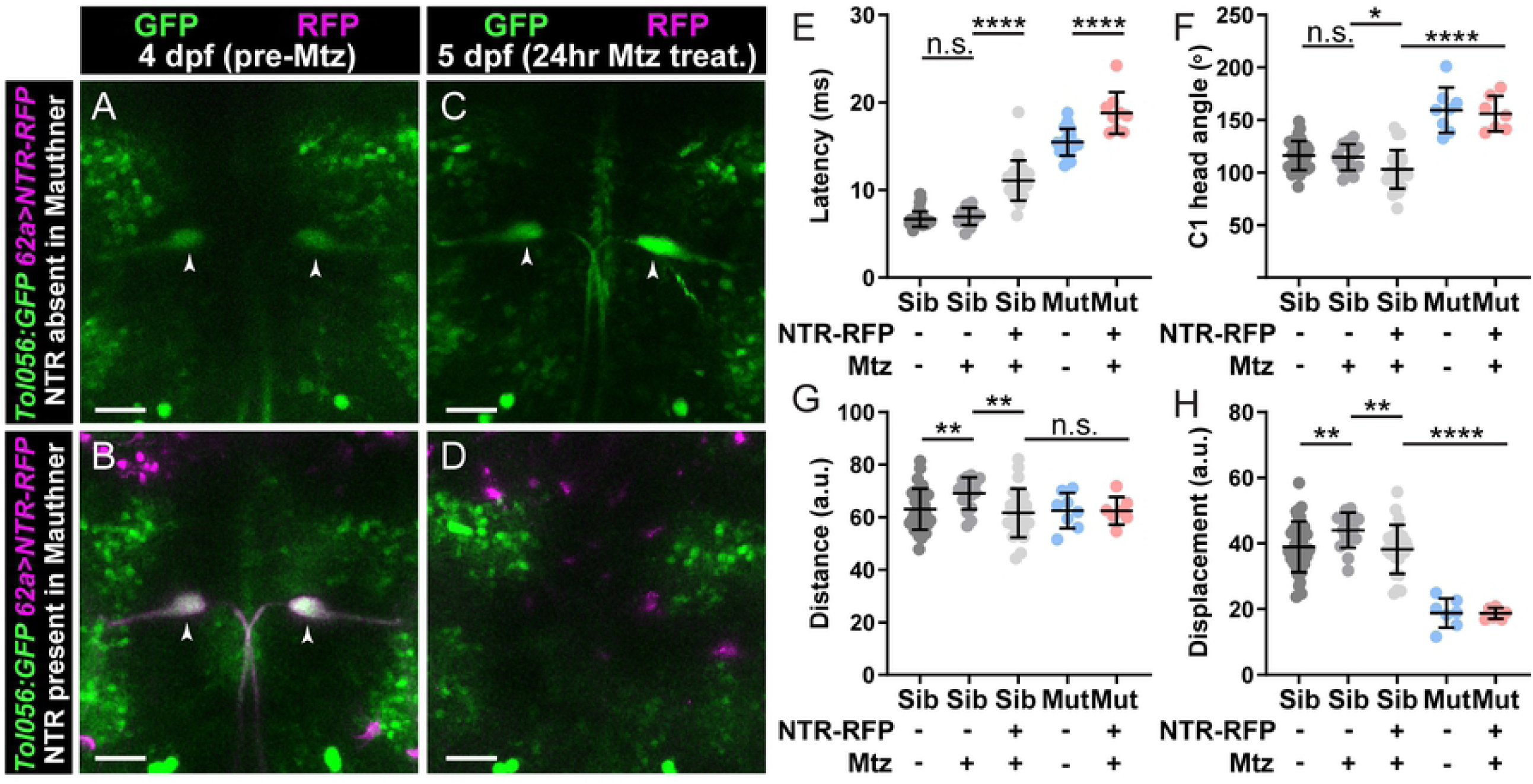
Kv1.1 functions outside the Mauthner command neuron to control swim movement magnitude. (A-D) *kcna1a* mutants expressing *Tol056* (GFP in Mauthner cells; green), *62a:Gal4* (Mauthner specific) and *UAS:NTR-RFP* (magenta). 62a>NTR-RFP is mosaic so some fish do not display expression in Mauthner cells (A) while others do (B). Before treatment with metronidazole (Mtz) at 4 dpf, all fish have GFP+ Mauthner cells (A,B). After 24 hr Mtz treatment, Mauthner cells are present in fish without initial NTR-RFP Mauthner expression (C) and absent in fish that displayed initial Mauthner NTR-RFP expression (D). (E-H) Kinematic parameters of the acoustic startle response in *kcna1a* sibling and mutant larvae with or without NTR-RFP expression in Mauthners and with or without Mtz treatment. Each point is the average response over ten trials for an individual larva. n≥9 larvae, *p=0.02, **p<0.01, ***p<0.0001. Scale bars=50 µM.

We next tested whether *kcna1a* acts in the spinal cord to regulate startle magnitude. We first determined whether *kcna1a* mutants exhibit any abnormal swim movements in the absence of external stimuli. *kcna1a* mutants execute normal swimming behaviors, including slow scoots and turns (Fig 6A,B), and when compared with their siblings, initiate a similar number of spontaneous movements (Fig 6G). However, after *kcna1a* mutant larvae begin a scoot maneuver, they sometimes undergo a series of rapid, high amplitude turns, similar to the behavior observed in response to acoustic stimuli (Fig 6C, Video S6). This exaggerated “thrashing” movement is not observed in *kcna1a* siblings (n=17 larvae, >800 responses), while *kcna1a* mutant larvae exhibit this behavior 22% of the time during spontaneous swim movements (Fig 6H). Thus, *kcna1a* regulates the magnitude of both acoustic startle responses and spontaneous swim movements.

**Fig 6.**
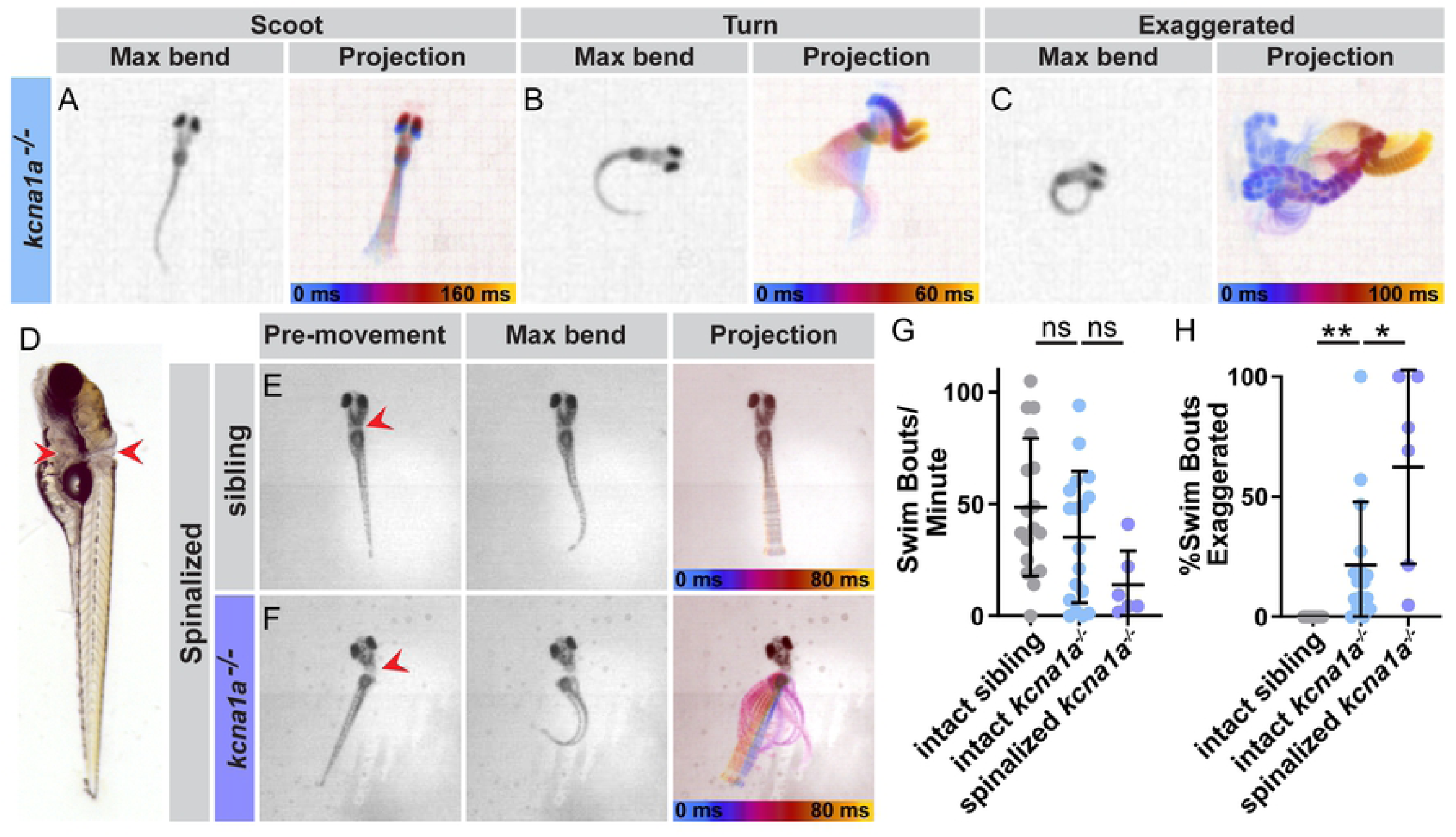
Kv1.1 functions in the spinal cord to control movement magnitude. (A-C) Examples of spontaneous swim movements performed by *kcna1a* mutant larvae. (D) Larvae were transected posterior to the brain to sever the spinal cord. Red arrows indicate the dorsal and ventral points of the cut. Some yolk was left intact to allow mounting of the head in agarose and imaging of the tails, which without the heads would lay on their sides. (E) Example of spinalized sibling and (F) spinalized *kcna1a* mutant spontaneous movement. (G) The number of spontaneous swim bouts initiated per minute for intact *kcna1a* siblings and mutants and spinalized *kcna1a* mutant larvae. (H) Percent of spontaneous swim bouts that are abnormal exaggerated movements with high amplitude bends. Each point is an individual fish. n≥6 fish. **p<0.0001, *p=0.03

Finally, to address whether Kv1.1 acts primarily in spinal circuits to drive movement, we removed brain input to the body by severing the spinal cord. Zebrafish larvae can be “spinalized” by severing the head from the trunk and leaving only the spine intact. These spinalized larvae are unable to perform acoustic startle responses but are still capable of initiating spontaneous movement [59], though the rate of movement is reduced. We therefore transected the spinal cords of *kcna1a* sibling and mutant larvae (Fig 6D) and observed their spontaneous movement. Compared to intact *kcna1a* mutant larvae, spinalized *kcna1a* mutant larvae continue to exhibit exaggerated swim bouts not observed in spinalized siblings (Fig 6E-H, Video S7). Combined, these results strongly suggest that Kv1.1 acts to control swim movements within spinal locomotor circuits.

## Discussion

Movement initiation, magnitude, and duration all depend on neural circuits acting in a coordinated manner to drive muscle contraction. Here, we describe mutations in eight genes identified in a forward genetic screen in larval zebrafish that enable and ensure coordinated movements. The driving force for carrying out this screen was to discover, in an unbiased way, genes that are required for regulating the acoustic startle response. With this unbiased approach, we predicted we would identify both genes that are already implicated in locomotion and genes without a prior known role in locomotion. This is in fact what we found. All of the eight genes we identified in our screen are associated with human disorders, and six are associated with motor disorders (Table 1). For example, *NEB* mutations in humans are associated with nemaline myopathy, a congenital muscle disease characterized by muscle weakness [31]. We identified *neb* mutants based on their weak response to an acoustic stimulus, and a previously characterized zebrafish *neb* mutant was shown to have defects in sarcomere assembly [32], similar to human patients with nemaline myopathy. Human mutations in *SLC5A7* are associated with congenital myasthenic syndrome [60], which is characterized by muscle weakness and fatigue. *slc5a7a* mutants from our screen display a weak acoustic startle response similar to *neb* mutants. While Neb protein acts within the muscle, Slc5a7 acts within motor neurons at the NMJ. Therefore, while the mutants have a similar phenotype, suggesting a similar site of action, the proteins act in different cell types. This underscores the importance of understanding the normal biological function of a protein when trying to understand the underlying cause of a genetic disorder.

Even when the function of a protein is known, the protein may act in many different neurons or cell types to regulate movement. For example, Kv1.1 is broadly expressed in the zebrafish nervous system ([54] and our work) and in the mammalian nervous system [53]. In humans, heterozygous disease alleles of the Kv1.1-encoding gene *KCNA1* have been shown to be causative for episodic ataxia (EA) Type 1 [61,62]. EA patients display frequent attacks of incoordination that are usually preceded by stress or startle. This phenotype is attributed to dysfunction in the cerebellum and peripheral nerves [63]. Work in mice heterozygous for a human *KCNA1* disease allele has demonstrated a cerebellar defect of action potential broadening in basket cells [64], and studies in patients with EA have demonstrated increased nerve excitability when the median nerve of the wrist was stimulated [65]. Thus, Kv1.1 acts at multiple points in the mammalian nervous system to regulate nerve activity.

In this study, we uncovered a brain-independent role for Kv1.1 in regulating swim movements in larval zebrafish. Spinalizing *kcna1a* mutant larvae fails to rescue motor dysfunction, suggesting that Kv1.1 plays a critical role in spinal motor circuits. Is Kv1.1 acting in interneurons, motor neurons, or both to drive movement? In *Drosophila*, there is a single Kv1 channel, Shaker, which was the first potassium channel ever cloned [66-68]. *Shaker* mutants exhibit aberrant movements and display broadened action potentials at NMJs [69]. In mice, Kv1.1 plays an important role in preventing spontaneous activity of motor neurons [70,71]. Kv1.1 is broadly expressed in the zebrafish larval spinal cord (Fig 4C), including in regions containing motor axons. However, the motor defect observed in zebrafish *kcna1a* mutants is distinct from other mutants that exhibit prolonged neurotransmitter release at the NMJ. For example, *twister* mutants harbor a gain-of-function mutation in the α-subunit of the muscle nicotinic acetylcholine receptor, resulting in prolonged synaptic transmission at the NMJ [72]. While wild type larvae respond to touch by performing rapid alternating swim movements to escape, *twister* mutants demonstrate loss of alternating left/right muscle contractions without productive forward movement [73]. In contrast, while we observe exaggerated turn angles in *kcna1a* mutants, swim movements still alternate from left to right to produce productive swimming. Considering these differences in phenotypes, it is unlikely that the zebrafish *kcna1a* mutant phenotype is due solely to increased motor neurons activity.

Another potential site of action for Kv1.1 in the spinal cord is in activating interneurons that excite motor neurons. Classes of locomotor interneurons are relatively conserved between fish and mammals [74]. One excitatory class is the V2a glutamatergic Chx10+ interneurons. In larval zebrafish, V2a interneurons are required for producing a normal acoustic startle response. When V2a interneurons are silenced, the C1 turn angle and tail beat frequency following the acoustic stimulus are reduced [75]. Recently, V2a interneurons in larval zebrafish were characterized as Type I or Type II, similar to characterizations in mice, based on morphological and electrophysiological properties. From this work, type II V2a interneurons are predicted to regulate amplitude control [76]. Loss of Kv1.1 in type II V2a interneurons could result in hyperexcitability or broadening of the action potential, leading to increased recruitment or activation of motor neurons. This could explain the movement amplitude defects observed in *kcna1a* mutant larvae. From single nucleus RNA-seq in the mouse spinal cord, *Kcna1* is enriched in V2a interneurons [77]. V2a interneurons are critical in mice to coordinate left-right alternating movement, particularly at high speeds [78]. Kv1.1 may play a critical role in zebrafish and mammals in regulating locomotion through limiting excitability of V2a interneurons.

While localization to different cell types provides specificity to Kv1.1’s function, post-translational modifications critically regulate its activity. We have previously shown that the palmitoyltransferase Hip14 palmitoylates Kv1.1, and both Hip14 and Kv1.1 are essential for zebrafish larvae to habituate to repeated acoustic stimuli [24]. Here, we demonstrate a role for glycosylation in regulating Kv1.1 function. In cell culture, glycosylation of Kv1.1 is required for proper channel function [49-51]. Interestingly, in these studies, glycosylation is not required for channel localization to the membrane. Here, we find that in *dolk* mutants, where Kv1.1 is predicted to not be glycosylated, localization of Kv1.1 is drastically disrupted. This effect may be directly related to the loss of glycosylation or may be an indirect effect caused by reduced glycosylation of trafficking proteins. For example, Leucine-rich glioma-inactivated 1 (LGI1) is a glycoprotein that promotes axonal localization of Kv1.1 in hippocampal cells [79]. Whether the effect is direct or indirect, loss of Dolk clearly drastically affects Kv1.1 localization.

How does loss of a ubiquitous protein cause nervous system specific defects? Protein glycosylation is required in all cell types, and Dolk is a critical part of the protein glycosylation pathway. *dolk* mRNA has been detected in all human tissues assayed, with the highest expression observed in the brain [80]. Patients with *dolk* disease alleles display a range of phenotypes. In some patients, homozygous loss of function mutations in *dolk* cause metabolic and heart defects that are often lethal early in life [80,81]. Patients with similar homozygous, loss of function *dolk* mutations have also been identified that only present with neurological conditions, including seizures, at the time of diagnosis in childhood [82]. A *dolk* variant has also recently been reported as a potential risk gene in a patient with autism spectrum disorder [83]. The neurological symptoms observed in patients with *dolk* mutations underscore the importance of protein glycosylation in the nervous system [84]. In fact, defects in the nervous system are a predominant feature of congenital disorders of glycosylation [85]. In zebrafish *dolk* mutants, we do not observe any obvious morphological defects by 5 dpf except for failure to inflate the swim bladder, which is common in mutants with swim defects [11]. We have not recovered homozygous *dolk* mutant adults. Post-developmental lethality could be due to a failure to eat since zebrafish larvae must perform coordinated swim movements to hunt and eat their food. Lethality could also be due to a requirement of *dolk* in other organ systems. The behavioral phenotype in these mutants, however, is very clear. While Kv1.1 is likely not the only disrupted protein in *dolk* mutants, the similarity in behavioral phenotypes of *kcna1a* and *dolk* mutants suggests Kv1.1 is a critical protein affected by loss of *dolk*. It would be interesting to investigate whether similar dysregulation of Kv1.1 occurs in patients with neurological phenotypes arising from *dolk* mutations.

Together, this work demonstrates the utility of zebrafish for understanding genetic mechanisms driving regulation of locomotion. While the neural circuits driving movement are not identical among species, many of the genes required are conserved. This is especially apparent from the number of genes identified from our screen that are associated with locomotor disorders in humans. In addition, while startle behaviors differ between species, the behavioral phenotypes associated with gene mutations are in some instances remarkably similar. For example, mice with *kcna1* mutations are hypersensitive to acoustic stimuli, display increased response latency during the acoustic startle response, and demonstrate increased amplitude of the response [86], similar to the acoustic startle phenotypes we observe in *kcna1a* mutant zebrafish larvae ([23] and this study). Further understanding of how loss of critical genes affects locomotor behaviors will shape our models of how molecular pathways act in behavioral circuits to drive movement.

## Supporting figure legends

**Fig S1. The high affinity choline transporter Slc5a7a is required in motor neurons for the startle response**. (A) Gene structure for *slc5a7a*, with the nonsense mutation from the screen (allele *p417*) noted. (B) Protein structure for Slc5a7a on the plasma membrane, with the predicted amino acid change from the screen mutation indicated. In contrast to siblings (C-G), *slc5a7a* mutants (H-L) display a weak startle response, with reduced bend and counter bend angles, as well as uncoordinated movement. Blue bar=latency in G,L. (M) *slc5a7a-/-; Tg(hb9;slc5a7a, hb9:mKate)* larvae express mKate and *slc5a7a* in motor neurons. mKate is expressed from a second *hb9* promoter to allow visualization of expressing neurons without disrupting protein function. Bracket indicates motor column and arrow indicates motor nerve axons. Scale bar=50 µM. (N-P) Kinematic parameters of the acoustic startle response in *slc5a7a* siblings and mutants with or without *hb9:slc5a7a* transgene. Each point represents an average of ten trials for an individual fish. n≥15 larvae, *p=0.002, **p<0.0001.

**Fig S2. Electrophysiological properties of the Mauthner cell are unaffected in *kcna1a* mutant larvae**.

(A) Averaged measurements of V_resting_, V_threshold_, Rheobase, and R_in_ in *kcna1a* sibling and mutant larvae. n.s.=not significant. (B) Representative mixed synaptic responses on the Mauthner cell evoked by electrical stimulation of auditory afferents (club endings) in *kcna1a* sibling (left) and mutant (right) larvae (responses represent the average of at least 10 single traces). The maximal amplitude of the electrical component (C), chemical component (D), and paired-pulse ratio of the chemical component (E) of the mixed synaptic response in *kcna1a* sibling or mutant Mauthner cells are not significantly different. n=7 *kcna1a* sibling larvae, n=6 *kcna1a* mutant larvae.

## Supporting video legends

**S1 Video. Acoustic startle response of 5 dpf wild type larva**. Stimulus is given at the start of the video and 90 ms of the response are shown. Video is slowed down 50x.

**S2 Video. Acoustic startle response of 5 dpf *ryr1b* mutant larva**. Stimulus is given at the start of the video and 90 ms of the response are shown. Video is slowed down 50x.

**S3 Video. Acoustic startle response of 5 dpf *prdm12b* mutant larva**. Stimulus is given at the start of the video and 90 ms of the response are shown. Video is slowed down 50x.

**S4 Video. Acoustic startle response of 5 dpf *slc5a7a* mutant larva**. Stimulus is given at the start of the video and 90 ms of the response are shown. Video is slowed down 50x.

**S5 Video. Acoustic startle responses of 6 dpf *kcna1a* mutant larvae with or without intact Mauthner neurons**. Larva with intact Mauthner neurons is on the left, and larva with ablated Mauthner neurons is on the right. Stimulus is given at the start of the video and 90 ms of the response are shown. Video is slowed down 50x.

**S6 Video. Examples of spontaneous movements by a *kcna1a* mutant larva**. Example of normal scoot, normal turn, and abnormal exaggerated movement are shown. Time labels refer to each individual movement type. Video is slowed down 50x.

**S7 Video. Abnormal movements of a spinalized *kcna1a* mutant larva**. Larva has had the spinal cord severed and the head is mounted in agarose. Three examples of spontaneous abnormal exaggerated movements are shown. Time labels refer to each individual movement type. Video is slowed down 50x.

## Methods

### Zebrafish husbandry and lines

All animal protocols were approved by the University of Pennsylvania Institutional Animal Care and Use Committee (IACUC). Embryos were raised at 29°C on a 14-h:10-h light:dark cycle in E3 media. *Tol-056* is from [22]. *hspGFF62A* (*62a:Gal4*) is from [87]. *UAS:NTR-RFP* is from [88].

### Behavior testing

Behavioral experiments were performed using 5 dpf larvae (except for Mauthner ablation experiments performed at 6 dpf) and analyzed using FLOTE software as described previously [16]. Larvae were tested in groups in a 6×6 laser cut grid. The acoustic stimuli used for acoustic startle experiments were 25.9 dB based on larval response rates and previous measurements of the speaker [25]. 10 stimuli were given with a 20 sec interstimulus interval. Before FLOTE tracking and analysis, ImageJ was used to remove background from videos to prevent erroneous measurements from the outline of the wells. Latency measurements in Table 1 were determined manually, as many mutants don’t have swim bladders and lay on their sides and FLOTE did not always reliably calculate the movement latency. Angle measurements by FLOTE that were incorrect due to larvae laying on their sides and turning in the wrong plane were discarded. For measurements of spontaneous movement, only larvae that moved in a three-minute period were counted. Videos of acoustic startle response were acquired at 1000 fps, and videos of spontaneous movement were acquired at 500 fps. Images in figures have background removed, except for the spinalized larvae, and some images were mirrored to show an initial rightward turn for ease of comparison between genotypes.

### Mutagenesis, WGS/WES, and transgenesis

ENU mutagenesis was performed using TLF and WIK as described previously [23].

Cloning of *p181* is described in [24].

Alleles *p415, p417*, and *p420* were cloned using WGS as described previously [25]. Alleles *p413, p414, p416*, and *p419* were cloned using WES as described previously [28]. For SNP analysis, we presumed our sibling pools (wild type behavior) consisted of 66% heterozygous (+/-) larvae and 33% (+/+) larvae, which results in a 1:2 ratio of mutant:wild type chromosomes in the analysis. To identify causative mutations and maintain mutant lines, mutations were genotyped using proprietary KASP primers and reagents (LGC Genomics). To confirm the splice site mutation in *ryr1b*^*p414*^, a forward primer was designed in exon 4 and a reverse primer was designed in exon 5. PCR performed on cDNA from *ryr1b* homozygous mutant larvae resulted in a larger product (245 bp), caused by retention of an intron, than the product from wild type larvae (153 bp).

To generate *dolk*^*p421*^, a CRISPR-Cas9 Alt-R gRNA (IDT) was designed using CHOPCHOP [89]. gRNA was combined with Cas9 protein (PacBio) and injected into one-cell stage embryos. F1 progeny from injected embryos were screened for mutations by sequencing the region flanking the gRNA site and using the ICE Analysis tool (v2) (Synthego) to identify Cas9 induced indels. Mosaic injected carriers were outcrossed to establish stable lines.

Unless noted otherwise, homozygous mutants were identified by abnormal kinematic behavior and siblings (heterozygous carriers and wild type) were designated as larvae performing normal startle responses. All phenotypes described in the paper display 100% penetrance, based on sequencing of at least twenty larvae displaying the mutant phenotype and at least twenty larvae displaying the wild type phenotype in presumed siblings. We do not observe phenotypes in heterozygous carrier larvae in any of the mutants described. Unless otherwise noted, all mutant alleles discussed in the text are from the forward genetic screen.

To generate *Tg(hb9:slc5a7a,hb9:mKate)*^*p418*^, *slc5a7a* was cloned from 5 dpf larval cDNA into PENTR-D-TOPO (Invitrogen). Gateway cloning was performed with a double *hb9* promoter pDest plasmid that includes I-SceI sites [90] to make *hb9:slc5a7a,hb9:mKate*. The construct and I-SceI were injected, as described previously [91], into one-cell stage embryos from an outcross of *slc5a7a*^*p417/+*^. Successful integration was monitored by screening for mKate in the spinal cord. In Figure S2, genotype was determined by screening for mKate expression and genotyping for *mKate* and *p417* mutation.

### Immunohistochemistry

6-7 dpf larvae were sorted by behavioral phenotype. Larvae were fixed in 2% trichloroacetic acid (TCA) in 1xPBS for three hours. Larvae were washed with PBS-0.25% Triton, blocked in 2% Normal Goat Serum, 1% BSA, 1% DMSO in PBT, and incubated with 1/200 α-Kv1.1 (rabbit, Millipore AB5174) and 1/50 α-3A10 (mouse, Developmental Studies Hybridoma Bank) in block solution for two days at 4°C. Following washes in PBT, larvae were incubated in secondary antibody overnight (1/400 anti-mouse 488, anti-rabbit 633; Millipore) at 4°C. After washing, larvae were dehydrated progressively in 25%, 50%, and 75% glycerol in PBS. Larvae were peeled to separate the spinal cord and brain and mounted ventral side towards the coverslip. Imaging was performed using a Zeiss LSM 880 confocal microscope.

### Electrophysiology

To perform electrophysiological recordings from *kcna1a* siblings and mutants, larvae (5– 8 dpf from an incross of *kcna1a*^*p181/+*^;*Tol-056*) were anesthetized using a 0.03% Tricaine solution pH adjusted to 7.4 with NaHCO_3_. Larvae were then paralyzed using d-tubocurarine (10 μM, Sigma) in external solution in mM: 134 NaCl, 2.9 KCl, 2.1 CaCl_2_, 1.2 MgCl_2_, 10 HEPES, 10 Glucose, pH adjusted to 7.8 with NaOH [49]. Then, larvae were transferred dorsal-side down to a Sylgard-coated small culture dish (FluoroDish, WPI) and immobilized with tungsten pins. The brain was exposed ventrally as described in [17]. Following surgery, the petri dish containing the larva was placed on stage of an Axio Examiner upright microscope (Carl Zeiss AG) adapted for electrophysiology recordings. Larvae were superfused with external solution throughout the recording session. Mauthner cells were identified by far-red DIC optics and GFP fluorescence. The patch pipette (3-4 MΩ) was filled with internal solution in mM: 105 K-Methanesulfonate, 10 HEPES, 10 EGTA, 2 MgCl_2_, 2 CaCl_2_, 4 Na_2_ATP, 0.4 Tris-GTP, 10 K_2_-Phosphocreatine, 25 mannitol, pH adjusted to 7.2 with KOH. Whole-cell recordings under current-clamp configuration were performed using a Multiclamp 700B patch-clamp amplifier connected to a Digidata 1440A (Molecular Devices) digitizer. The liquid junction potential was estimated in -16 mV using Clampex 10.6 (Molecular Devices). The rheobase, defined as the minimum amount of positive current needed to elicit an action potential, was determined by delivering a 20 ms current pulse. Voltage threshold was measured as the membrane potential value at which the depolarizing-current step elicits an action potential. The input resistance was estimated using the voltage deflection caused by a hyperpolarizing-current step of -1 nA and 20 ms duration, followed by derivation of resistance with Ohm’s law. To activate the auditory afferents to the Mauthner cell, a “theta” septated glass pipette filled with external solution was used as bipolar electrode. The bipolar electrode was placed at the posterior macula of the ear, where the dendritic processes of auditory afferents contact the hair cells [57].

### Mauthner cell ablation

Larvae (obtained from a cross of *kcna1a*^*p181*^*/+;Tol-056/+;UAS:NTR-RFP/+* and *kcna1a*^*p181*^*/+;62a:Gal4/+*) were sorted at 60 hpf for GFP+ (*Tol-056*) and NTR-RFP+ on a fluorescent dissecting scope. At 4 dpf, larvae were behavior tested and sorted into mutants and siblings. Because *62a>NTR-RFP* is mosaic, larvae expressing NTR-RFP in the brain were then mounted in agarose and screened on a compound epifluorescent scope for specific expression of NTR-RFP+ in the Mauthner cells. Only fish where both or neither Mauthner cells displayed expression were used. At 104 hpf, larvae were treated with 10mM Metronidazole/Mtz (Sigma-Aldrich) in 0.2% DMSO in E3. After 24 hours, Mtz was washed out and larvae were mounted and screened again to confirm ablation. At 144 hpf, larvae were behavior tested. Representative images of live fish in Figure 5 were obtained from a Zeiss LSM 880 confocal microscope.

### Larvae spinalization

5 dpf larvae were anesthetized with Tricaine (Sigma-Aldrich) in Hank’s balanced salt solution supplemented with 1x Glutamax and 1x sodium pyruvate. A sliver from a double edge razor was used to cut through the spinal cord and musculature through into the yolk, posterior to the brain, roughly at the halfway point of the swim bladder. Larvae were mounted in 2% low melt agarose in supplemented Hank’s and agarose covering the tail was removed. Larvae were allowed to recover 30 min-2 hours in Hank’s solution without Tricaine, until spontaneous movement was observed.

## Statistics

Statistical analyses were performed using GraphPad Prism. Pairwise comparisons were performed using non-parametric Mann-Whitney tests. All plots display mean and standard deviation.

## Acknowledgements

The authors would like to thank Mary Mullins, Shannon Fisher, Bill Vought, Paula Roy, Hannah Bell, and Julianne Skinner for help with the genetic screen; Harold Burgess for *UAS:NTR-RFP*; the UPenn Cell & Developmental Biology Microscopy Core, UPenn Next-Generation Sequencing Core (WGS), and Duke Sequencing and Genomic Technologies Shared Resource (WES) for equipment and sequencing; Katharina Hayer for design of WGS pipeline; and members of the Granato lab for feedback regarding the manuscript.

## Author contributions

JHM, JCN, KCM, RAJ, MAW, and MG designed the research; JHM, JCN, KCM, JH, RAJ, and MAW performed research and analyzed data; JHM and MG wrote the paper.

